# How sleeping minds decide: state-specific reconfigurations of lexical decision-making

**DOI:** 10.1101/2025.03.10.642374

**Authors:** Tao Xia, Chuan-Peng Hu, Basak Türker, Esteban Munoz Musat, Lionel Naccache, Isabelle Arnulf, Delphine Oudiette, Xiaoqing Hu

## Abstract

Decision-making is a core cognitive function that enables adaptive behavior across diverse contexts. While extensively studied in wakefulness, its persistence and reconfiguration across sleep states remain poorly understood. Here, we use computational modeling to examine lexical decision-making across wakefulness, N1 sleep, and lucid REM sleep in both healthy participants (HP) and participants with narcolepsy (NP). Using facial electromyography (EMG) to capture real-time behavioral responses to spoken words and pseudowords during sleep, we quantify how decision-making strategies adapt under different sleep and consciousness states. Our findings reveal two key insights. First, decision-making mechanisms are dynamically reconfigured across sleep states. In N1 sleep, the advantage for word (vs. pseudoword) judgments is supported by faster sensory encoding and motor preparation, combined with efficient evidence accumulation. In contrast, in lucid REM sleep, the word advantage is driven exclusively by enhanced evidence accumulation, while sensory encoding and motor preparation remain unchanged. Second, cross-state comparisons reveal distinct patterns of preservation and impairment. In N1 sleep, word judgment remains largely intact, whereas pseudoword judgment is significantly impaired, characterized by prolonged stimulus encoding, delayed motor preparation, and reduced evidence accumulation. In contrast, lucid REM sleep is marked by a global reduction in processing efficiency, reflected in slower evidence accumulation and elevated decision thresholds for both words and pseudowords. These results demonstrate that rather than being uniformly degraded, decision-making is dynamically reconfigured across sleep stages, reflecting adaptive neurocognitive strategies that sustain cognition in altered states of consciousness. By identifying state-specific computational mechanisms, this study provides new insights into the brain’s resilience and flexibility under changing cognitive and physiological conditions.

## Introduction

Decision-making is core for human cognition, supporting individuals to adaptively navigate complex environments. While decision-making during wakefulness has been extensively studied, how the brain supports this process during sleep remains poorly understood. Sleep is increasingly recognized as an active cognitive state, during which the brain continues to process sensory information, acquire and consolidate memories, and even engage in goal-directed behaviour (Andrillon et al., 2016; Arzi et al., 2012; Konkoly et al., 2021; Türker et al., 2023; Xia et al., 2024). Despite these intriguing findings (Konkoly et al., 2021; Türker et al., 2023), the computational mechanisms that enable decision-making to persist and adapt under the distinct sleep states remain unclear. Here, leveraging unique data involving lexical decision makings during sleep and lucid dreaming (Türker et al., 2023), we investigated how sleeping minds reconfigure its computational strategies across wakefulness and different sleep states. By uncovering the computational mechanisms underlying decision-making during sleep, our findings provide novel insights into how cognitive processes are dynamically restructured across altered states of consciousness.

Sleep is not a uniform state but a progression through distinct neural stages, each imposing unique constraints on cognitive processing. In N1 sleep, the transition from wakefulness is gradual; sensory processing remains partially intact, allowing recognition of familiar stimuli, but higher-order cognitive functions are reduced (Andrillon et al., 2016; Blume et al., 2017; Lacaux et al., 2024; Wislowska et al., 2022). As sleep deepens into N2 and N3, slow-wave activity increases, thalamocortical connectivity diminishes, and responsiveness to external stimuli further declines, with the brain prioritizing endogenous memory consolidation (Diekelmann & Born, 2010; Massimini, 2005; Strauss et al., 2015). In contrast, REM sleep— often referred to as "paradoxical sleep"—is characterized by wake-like cortical activity and vivid dreaming experiences (Brown et al., 2012; Hobson & Friston, 2012). A particularly intriguing phenomenon is lucid REM sleep, in which individuals become aware that they are dreaming and can even exert voluntary control over dream content (Filevich et al., 2015; Voss et al., 2014; Zerr et al., 2024). This state represents a unique hybrid of internally generated cognition and externally responsive awareness, bridging the gap between sleep and wakefulness and offering a powerful model for experimentally probing cognitive processes during sleep.

A recent study demonstrated that individuals in lucid REM sleep can perceive and respond to questions presented by an experimenter in real time, using predefined eye movements (EOG) or facial muscle contractions (EMG) (Konkoly et al., 2021). Moreover, EMG-based lexical decision tasks—where individuals distinguish words from pseudowords by contracting facial muscles—have revealed that lexical decision-making remains possible throughout different sleep states, including lucid REM sleep (Türker et al., 2023). These findings provide key evidence that the sleeping minds remain a remarkable ability not only to process external stimuli but also to engage in higher-level cognitive functions, such as lexical judgments. However, the computational mechanisms underlying decision-making across different sleep states remain poorly understood, requiring further investigation.

To address this question, we employed drift diffusion modelling (DDM), a computational framework widely used to quantify the cognitive processes underlying decision-making (Ratcliff et al., 2004, 2016). The DDM conceptualizes decision-making as the gradual accumulation of noisy sensory evidence until a decision threshold is reached (Myers et al., 2022). Key parameters include the drift rate: how efficiently evidence is accumulated; the non-decision time: the time needed for sensory encoding, motor preparation, and other non-decisional processes; and the decision threshold: the amount of evidence required to make a decision, reflecting the trade-off between speed and accuracy. Prior studies have shown that differences in lexical decision-making—such as distinguishing words from pseudowords— are primarily driven by drift rate and non-decision times (Donkin et al., 2009; Ratcliff et al., 2004). However, to account for potential variations in decision-making strategies across wakefulness and sleep, we also included the decision threshold (a) in our model. Adjustments to the decision threshold are particularly important in contexts like sleep, where individuals must navigate the trade-off between speed and accuracy (Ratcliff et al., 2004; Ratcliff & McKoon, 2008). By applying DDM to EMG-measured muscle responses during a lexical decision task, we quantified how these parameters shift across wakefulness, light NREM sleep, and lucid REM sleep, revealing the computational trade-offs that sustain decision-making under altered sleep states.

To gain deeper insight into how decision-making adapts across sleep states, we studied participants with narcolepsy, a condition characterized by unstable sleep-wake transitions and frequent lucid dreams (Baird et al., 2019; Dodet et al., 2015; Mota-Rolim & Araujo, 2013). Because individuals with narcolepsy often experience heightened dream awareness and control, this condition provides a unique natural model for investigating decision-making in lucid REM sleep. Participants completed a lexical decision task during both wakefulness and sleep, allowing us to examine how cognitive processes adapt across altered states of consciousness. Our findings reveal that core cognitive components—such as non-decition times, evidence accumulation, and decision thresholds—are not simply degraded during sleep. Instead, they are dynamically reconfigured, highlighting the brain’s remarkable ability to sustain cognition despite shifting cognitive states.

## Results

We first investigated the computational mechanisms underlying lexical decision-making across wakefulness, N1, N2, non-lucid REM, and lucid REM sleep. Using DDM, we examined how these mechanisms adapt across sleep states to determine how sleep alters decision-making processes. Specifically, we assessed behavioural performance—measuring reaction times (RTs) and accuracy—as well as computational parameters derived from DDM: non-decision time (reflecting sensory encoding and motor preparation), drift rate (reflecting the efficiency of evidence accumulation), and decision threshold (reflecting decision caution). By comparing these parameters across states, we aimed to determine the extent to which lexical decision-making is preserved, impaired, or reconfigured during different sleep states.

### Lexical decision-making during wakefulness

To establish a baseline for decision-making processes, we first examined lexical decision-making during wakefulness. As expected, participants responded faster to words than pseudowords, reflecting more efficient linguistic processing.

Among healthy participants (HP), RTs were significantly faster for words compared to pseudowords (median*_diff_* = −0.171, 95% HDI [−0.221, −0.126], Figure 2A right), while accuracy differences were not significant (median*_diff_* = −0.116, 95% HDI [−0.485, 0.243], Figure 2A left). DDM revealed that the RT advantage for words was driven by shorter non-decision times (median*_diff_* = −0.106, 95% HDI [−0.149, −0.061], Figure 2C), reflecting efficient stimulus encoding and motor preparation. Drift rates (evidence accumulation) and decision thresholds (decision caution) were similar for words and pseudowords (all 95% HDIs overlapped with 0, Figure 2C, 2E).

**Figure 1.**
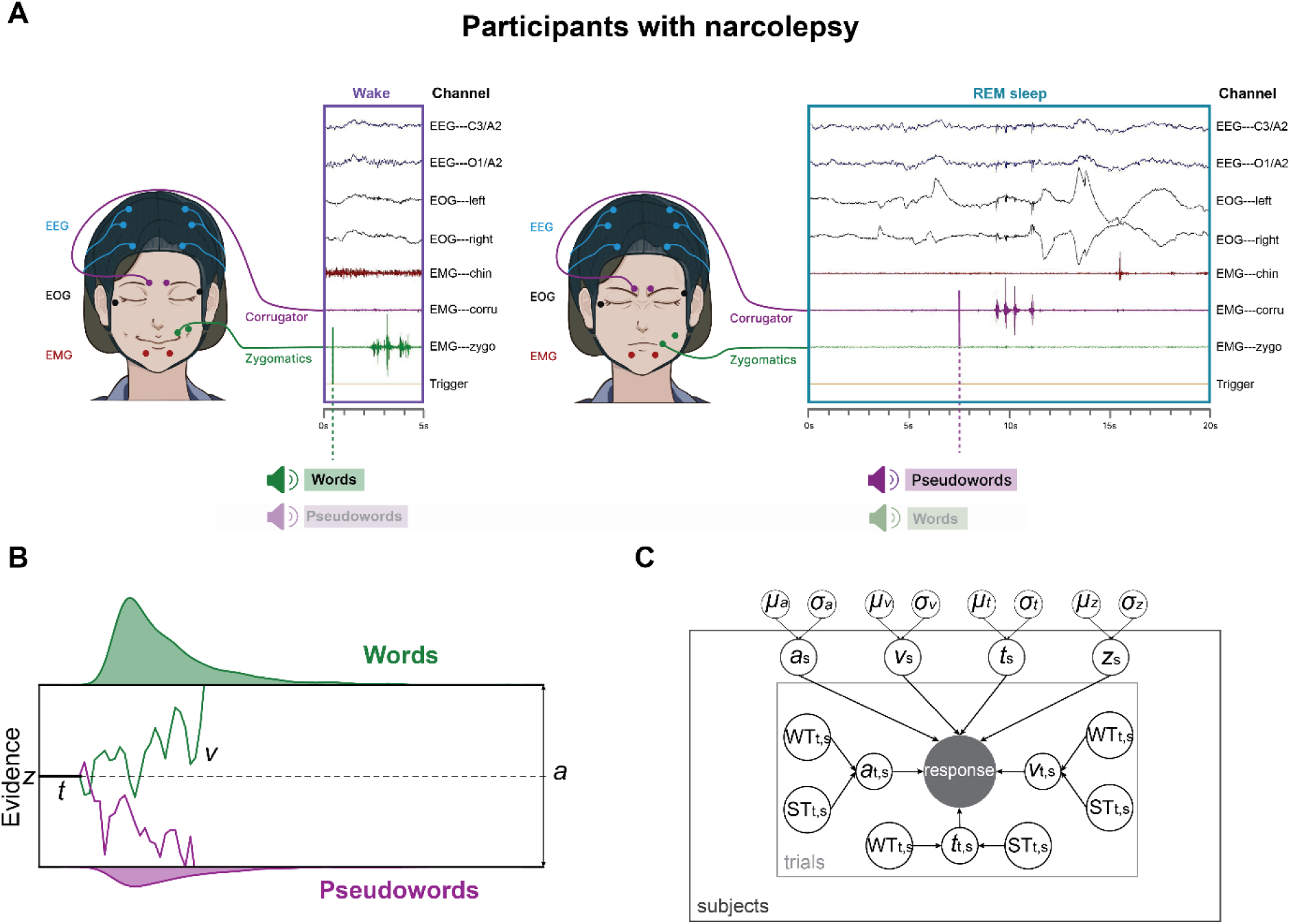
Experiment Design. A. Experimental Procedure. Participants performed a lexical decision task during daytime nap sessions, responding to spoken stimuli (words or pseudowords) with facial muscle contractions. Responses involved either frowning (corrugator muscle contractions) or smiling (zygomatic muscle contractions), with the muscle-response mapping counterbalanced across participants. Participants with narcolepsy (NP) completed five 20-minute naps, interspersed with 80-minute breaks, while healthy participants (HP) underwent a single 100-minute nap. Example EMG traces illustrate corrugator and zygomatic responses during wakefulness and REM sleep in NPs. Each participant was exposed to pseudorandomized auditory stimuli, ensuring no stimulus repetition across trials. B. Drift Diffusion Model (DDM) Schematic. The DDM decomposes decision-making into distinct cognitive components: the starting point (z), representing pre-decision bias; the drift rate (v), indicating the speed and quality of evidence accumulation; the decision threshold (a), reflecting the amount of evidence required to make a decision; and the non-decision time (t), encompassing processes unrelated to evidence accumulation (e.g., stimulus encoding, motor execution, and lexical access). The figure illustrates evidence accumulation over time, with green and purple traces representing correct and incorrect decisions, respectively. The model captures both trial-level response times and accuracy data. C. Hierarchical Bayesian HDDM Framework. A hierarchical Bayesian implementation of the DDM (HDDM) was used to estimate group- and participant-level parameters. Group-level parameters (mean, m, and variance, s) were estimated simultaneously with individual-level parameters (z, a, v, and t), accounting for trial-by-trial variations due to experimental factors (WT: word type; ST: sleep stages). At the trial level (T), parameters a, v, and t were modulated by word type (words vs. pseudowords) and sleep stages (wakefulness, N1, N2, REM, and lucid REM). Observed data (accuracy and reaction time) are represented as shaded circles, while group and individual parameters are shown as unshaded circles within the nested plate structure. This hierarchical approach improves parameter estimation by leveraging shared information across participants.

**Figure 2.**
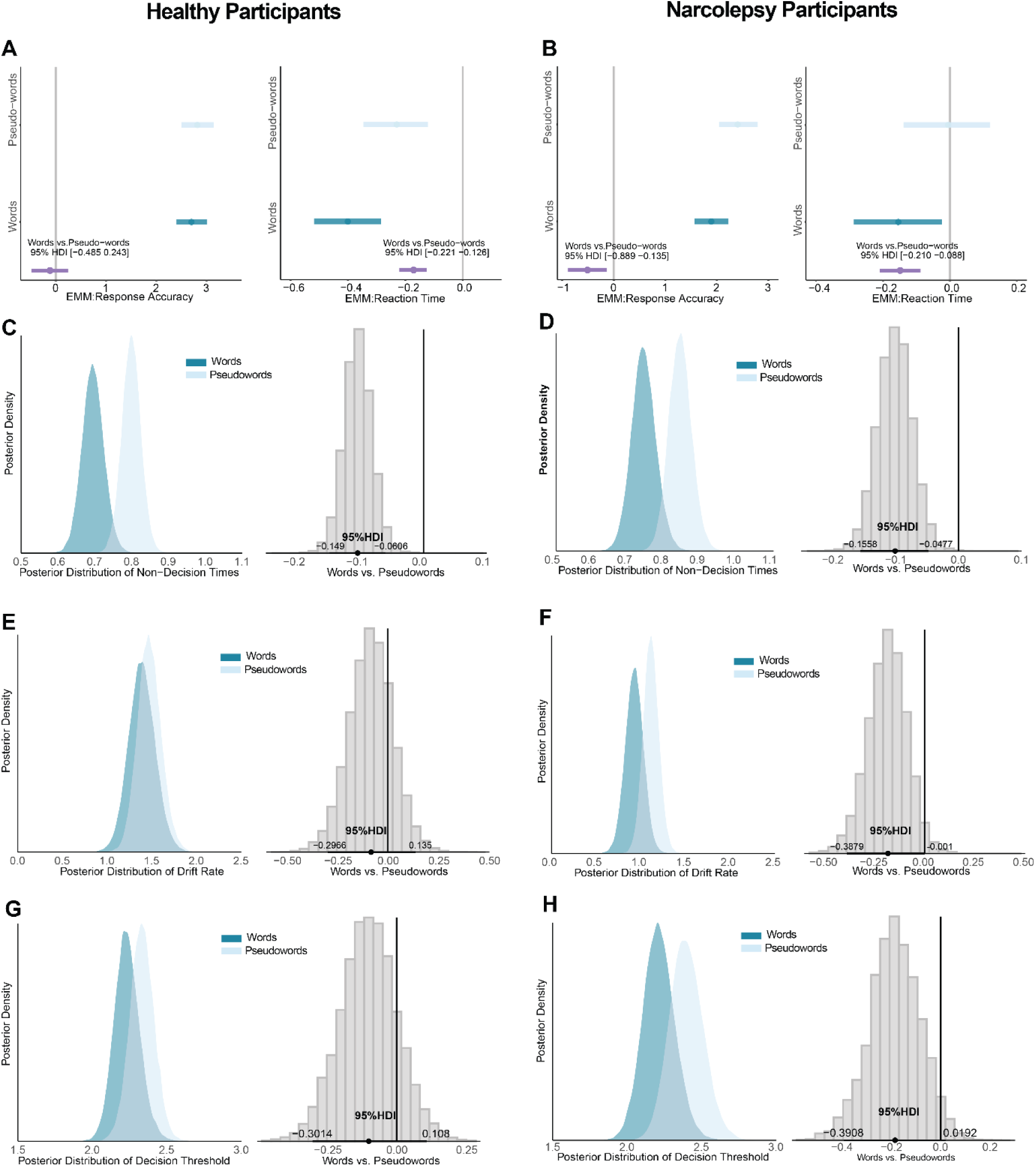
Lexical decision during wakefulness in both HP and NP. A, D. Behavioral Results: Response Accuracy and Reaction Time. Bayesian Linear Mixed Model (BLMM) results for lexical decisions in HP (left panels) and NP (right panels) during wakefulness. The x-axis represents the estimated mean for response accuracy (top) and reaction time (bottom). Light blue lines indicate the 95% Highest Density Interval (HDI) for pseudowords, and green lines indicate words. The purple line represents the posterior distribution of the contrast between words and pseudowords. If the purple line overlaps with 0 (gray vertical line), no significant difference is observed between words and pseudowords. Conversely, if the purple line does not overlap with 0, this indicates a significant difference between the two stimulus types. B, E. Drift Rate (v). Posterior distributions of drift rates for words (green) and pseudowords (light blue) during wakefulness in HP (left panels) and NP (right panels). The left panels show the posterior distributions for each word type, while the right panels display the posterior distribution of the contrast between words and pseudowords. The horizontal black lines represent the 95% HDI, and the vertical gray line (0) indicates the null hypothesis. Drift rate measures the speed and quality of evidence accumulation; a contrast excluding 0 indicates a significant difference in drift rates between words and pseudowords. C, F. Non-Decision Times (t). Posterior distributions of non-decision times for words (green) and pseudowords (light blue) in HP (left panels) and NP (right panels). Non-decision time reflects processes outside the evidence accumulation phase, such as sensory encoding and motor preparation. The left panels show the posterior distributions for each word type, while the right panels display the posterior distribution of the contrast between words and pseudowords. Horizontal black lines represent the 95% HDI, and vertical gray lines denote 0. If 0 is excluded from the 95% HDI, this indicates a significant difference in non-decision times between words and pseudowords.

In contrast, individual with narcolepsy (NP) exhibited abnormal lexical decision-making patterns. While RTs were faster for words than pseudowords (median*_diff_* = −0.149, 95% HDI [−0.210, −0.088], Figure 2B right), accuracy was lower for words (median*_diff_* = −0.512, 95% HDI [−0.889, −0.135], Figure 2B left). These deficits were reflected in faster non-decision times (median*_diff_* = −0.101, 95% HDI [−0.156, −0.048], Figure 2D) but significantly slower drift rates (median*_diff_* = −0.184, 95% HDI [−0.388, −0.001], Figure 2F) for words, suggesting impaired evidence accumulation. Decision thresholds were comparable across words and pseudowords (all 95% HDIs overlapped with 0, Figure 2G, 2H).

Thus, while HPs demonstrated efficient lexical decision-making during wakefulness, characterized by robust sensory encoding and motor preparation for words. NPs exhibited disrupted evidence accumulation, potentially reflecting cognitive impairments associated with narcolepsy (Naumann et al., 2006). Having established these baseline differences, we next examined how lexical decision-making is reconfigured across sleep states.

### State-specific mechanisms of lexical decision-making N1 and lucid REM sleep

Despite transitioning into sleep, participants retained the ability to make lexical decisions during both N1 sleep and lucid REM sleep, as indicated by faster and more accurate responses to words than pseudowords (see below). However, the mechanisms underlying this word advantage differed between these states.

In N1 sleep, RTs were significantly faster for words compared to pseudowords in both HPs (median*_diff_* = −0.390, 95% HDI [−0.575, −0.208], Figure S2A right) and NPs (median*_diff_* = - 0.160, 95% HDI [−0.265, −0.058], Figure 3A right). Accuracy was also higher for words than pseudowords in both groups (HP: median*_diff_* = 1.571, 95% HDI [0.479, 2.783], Figure S2A left; NP: median*_diff_* = 0.604, 95% HDI [0.107, 1.113], Figure 3A left). In lucid REM sleep, participants with narcolepsy similarly showed faster RTs (median*_diff_* = −0.169, 95% HDI [−0.281, −0.051], Figure 3B right) and higher accuracy (median*_diff_* = 0.674, 95% HDI [0.186, 1.161], Figure 3B left) for words compared to pseudowords. These findings indicate that lexical decision-making remains functional in both states.

**Figure 3.**
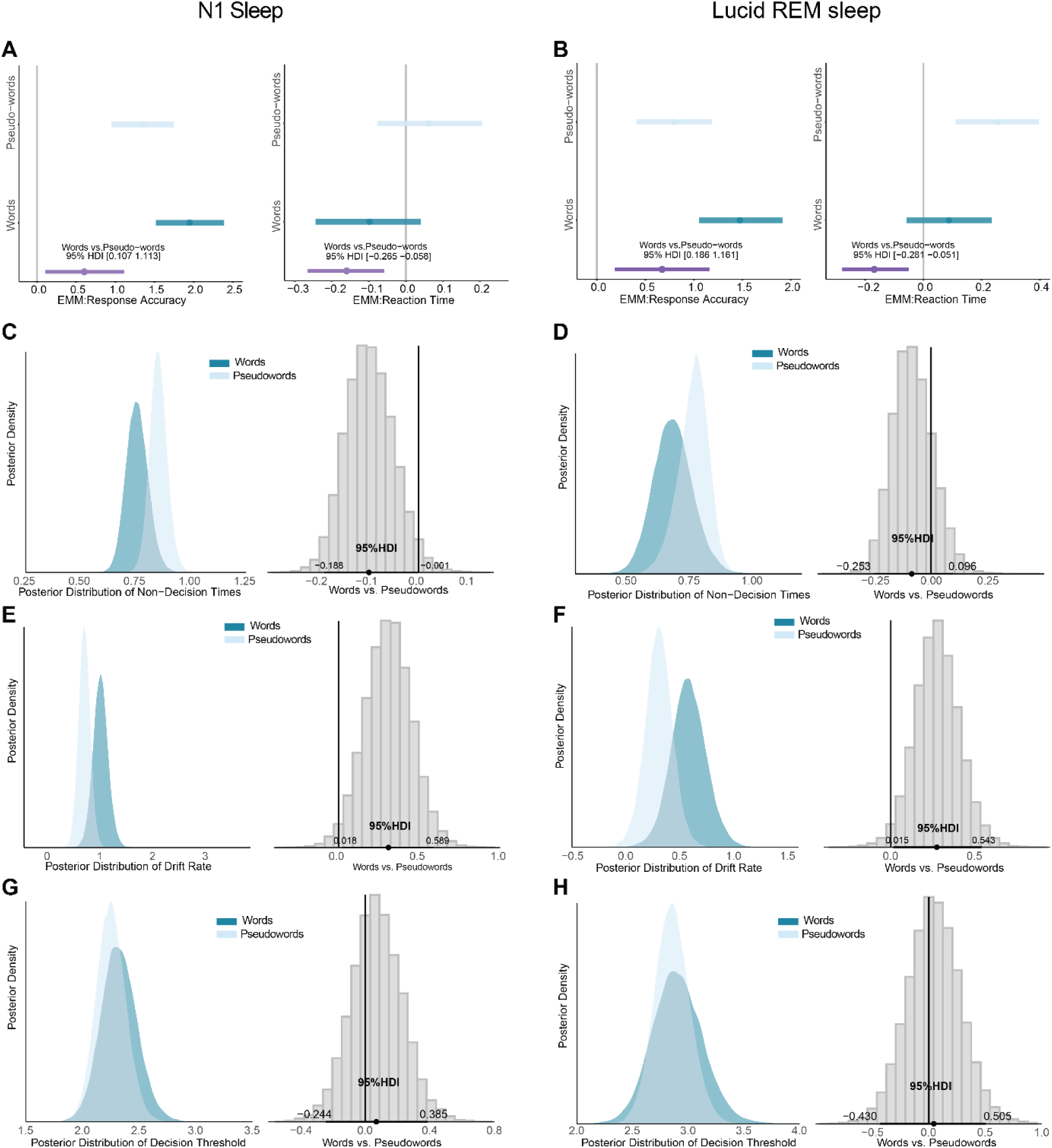
Mechanism of lexical decision during N1 and lucid REM sleep. (A, C) BLMM results for response accuracy and reaction time in lexical decisions during N1 sleep and lucid REM sleep in participants with narcolepsy. The N1 sleep results for healthy participants exhibited the same behavioral pattern and computational mechanisms as in participants with narcolepsy and are presented in Figure S2. The X-axis represents the estimated mean response accuracy or reaction time. The light blue line indicates the 95% Highest Density Interval (HDI) of the posterior probability for pseudowords, while the green line represents words. The purple line denotes the contrast between words and pseudowords. If the purple line overlaps with 0 (gray vertical line), there is no significant difference between words and pseudowords. If the purple line does not overlap with 0, it indicates a significant difference between words and pseudowords. (B, D) Posterior distributions of non-decision times, drift rate, and decision threshold for words and pseudowords during N1 and lucid REM sleep, along with their contrasts. The left panels show the fitted posterior distributions for words and pseudowords. The right-side histogram plots display the contrasts between words and pseudowords, with horizontal black lines representing the 95% HDI and vertical gray lines denoting 0. If 0 is not within the 95% HDI, the difference is considered statistically significant.

To uncover the mechanisms underlying lexical decisions during N1 and lucid REM sleep, we applied drift diffusion modelling. The results revealed distinct mechanisms supporting the word advantage in each state. In N1 sleep, the word advantage was driven by shorter non-decision times for words than pseudowords, reflecting more efficient sensory encoding and motor preparation in HPs (median*_diff_* = −0.213, 95% HDI [−0.367, −0.062], Figure S2B) and NPs (median*_diff_* = −0.098, 95% HDI [−0.188, −0.001], Figure 3C). Additionally, higher drift rate for words than pseudowords in HPs (median*_diff_* = 0.732, 95% HDI [0.105, 1.433], Figure S2C) and NPs (median*_diff_* = 0.308, 95% HDI [0.018, 0.589], Figure 3E), suggest enhanced evidence accumulation, facilitating more efficient lexical decision-making in both groups. Importantly, decision thresholds did not differ between words and pseudowords in both groups (all 95% HDIs overlapped with 0, Figure 3G, Figure S2E), indicating stable decision threshold across stimuli. These findings suggest that lexical decisions in N1 sleep were supported by preserved sensory encoding and motor preparation, along with efficient evidence accumulation.

In lucid REM sleep, the word advantage was primarily supported by higher drift rates for words than pseudowords (median*_diff_* = 0.274, 95% HDI [0.015, 0.543], Figure 3F), indicating selective improvements in evidence accumulation for words. Unlike N1 sleep, non-decision times and decision thresholds did not differ between words and pseudowords (all 95% HDIs overlapped with 0, Figure 3 DH), suggesting that sensory encoding and motor preparation, along with decision threshold were stable across stimulus types during lucid REM sleep. These findings suggest that lexical decisions in lucid REM sleep relied predominantly on selective improvements in evidence accumulation for words, rather than changes in non-decision times or decision caution.

Together, lexical decision-making in N1 and lucid REM sleep relied on distinct computational strategies. In N1 sleep, faster responses and higher accuracy were supported by both faster sensory encoding, motor preparation, and more efficient evidence accumulation. In lucid REM sleep, the word advantage was primarily driven by enhanced evidence accumulation, while sensory encoding, motor preparation, and decision caution remained unchanged.

### Absence of lexical decision-making in N2 and non-lucid REM sleep

In contrast to N1 and lucid REM sleep, participants were unable to distinguish between words and pseudowords in N2 and non-lucid REM sleep. There were no significant differences in RTs, accuracy, or any decision-making parameters (drift rates, non-decision times, or decision thresholds) between words and pseudowords (all 95% HDIs overlapped with 0, Figure S3). These findings indicate that lexical decision-making mechanisms are functionally absent in these deeper sleep states, likely reflecting reduced sensory processing and diminished cognitive engagement.

### Dynamic reconfiguration of lexical decision-making across wakefulness, N1 sleep, and lucid REM sleep

Having established that word judgments in N1 and lucid REM sleep rely on distinct computational mechanisms, we next examine how lexical decision-making adapts across wakefulness, N1 sleep, and lucid REM sleep. By integrating both behavioural performance and computational modelling, we identified gradual yet distinct changes in sensory encoding, motor preparation, evidence accumulation, and decision caution across these states, using wakefulness as a baseline for optimal performance.

### Wakefulness vs. N1 Sleep: robust word processing and impaired pseudoword processing

To examine how N1 sleep reconfigures decision-making processes for lexical decisions, we compared behavioural and computational results between wakefulness and N1 sleep in both HP and NP groups. The findings revealed a selective preservation of word processing, while pseudoword judgments were significantly impaired.

For words, RTs (median*_diff_* = 0.020, 95% HDI [−0.106, 0.148]), and accuracy (median*_diff_* = - 0.163, 95% HDI [−1.216, 0.765]) were comparable between wakefulness and N1 sleep, suggesting word judgment during N1 sleep remained stable. DDM confirmed that non-decision times, drift rate, and decision threshold for words did not significantly differ between wakefulness and N1 sleep (all 95% HDIs overlapped with 0, Figure 4A, 4B, 4C), suggesting that semantic networks supporting word judgment remained intact during N1 sleep.

**Figure 4.**
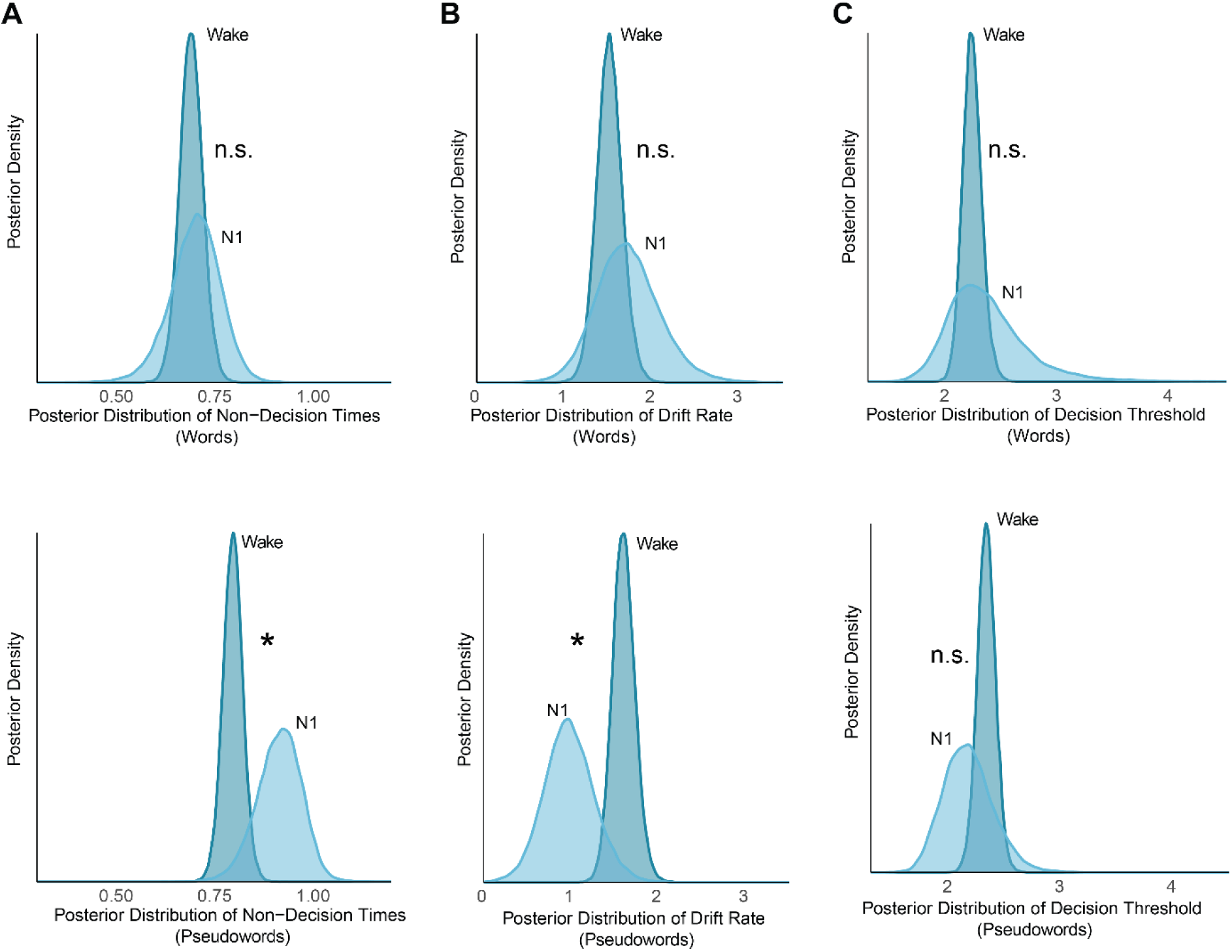
Robust word processing and impaired pseudoword processing during N1 sleep. Posterior distributions of decision parameters for words and pseudowords in the HP group, comparing wakefulness and N1 sleep. The NP group exhibited a similar pattern, with selective impairment in pseudoword processing during sleep (see Results). Panels depict (A) non-decision time, (B) drift rate, and (C) decision threshold. Asterisks (*) indicate significant differences, where the posterior distribution contrast between wakefulness and N1 sleep does not overlap with 0. "n.s." denotes non-significant differences, where the posterior distribution contrast overlaps with 0.

In contrast, pseudoword judgment was significantly impaired during N1 sleep. RTs were slower (median*_diff_* = −0.199, 95% HDI [−0.345, −0.057]), and accuracy was lower (median*_diff_* = 1.519, 95% HDI [0.823, 2.236]), reflecting greater difficulty in processing novel or ambiguous stimuli during N1 sleep. DDM revealed that this impairment was driven by slower non-decision times (median*_diff_* = −0.120, 95% HDI [−0.215, −0.009], Figure 4D) and lower drift rate (median*_diff_* = 0.630, 95% HDI [0.074, 1.115], Figure 4E), while the decision threshold remained unchanged between wakefulness and N1 sleep (95%HDI overlapped with 0, Figure 4F).

A similar pattern was observed in the NP group, where pseudoword processing exhibited a significantly slower drift rate compared to wakefulness (median*_diff_* = −0.468, 95% HDI [−0.688, −0.252]), while non-decision times and decision thresholds remained unchanged (all 95% HDIs overlapped with 0). However, as with HPs, word processing remained stable across wakefulness and N1 sleep, with no significant differences in non-decision times, drift rates, or decision thresholds (all 95% HDIs overlapped with 0).

These findings suggest that while lexical representations remain accessible during N1 sleep, decisions requiring phonological decoding and inhibitory control (e.g., pseudowords) are disproportionately affected, likely due to sleep-related reductions in cognitive resources. The similarity of this pattern across both groups supports the idea that familiar word judgment is preserved during N1 sleep, whereas processing novel or ambiguous stimuli becomes less efficient due to state-dependent cognitive constraints.

### Lucid REM sleep vs. wakefulness and N1 sleep: slower and less efficient lexical decisions

Building on these observations, we next examined how lucid REM sleep reconfigured decisional processing to perform lexical decision relative to wakefulness and N1 sleep. Unlike N1 sleep, where word processing remained largely intact, lucid REM sleep was associated with global reductions in processing efficiency.

Participants exhibited slower RTs across both stimulus types during lucid REM sleep. For words, RTs were significantly slower compared to wakefulness (median*_diff_* = 0.242, 95% HDI [0.141, 0.344]) and N1 sleep (median*_diff_* = 0.185, 95% HDI [0.075, 0.308]). A similar pattern was observed for pseudowords, with RTs significantly slower compared to wakefulness (median*_diff_* = 0.261, 95% HDI [0.161, 0.366]) and N1 sleep (median*_diff_* = 0.195, 95% HDI [0.080, 0.312]).

Despite the generalized slowing, accuracy patterns differed between stimulus types. Word accuracy remained stable, showing no significant differences compared to wakefulness (median*_diff_* = −0.418, 95% HDI [−0.916, 0.045]) or N1 sleep (median*_diff_* = −0.474, 95% HDI [−1.011, 0.088]). In contrast, pseudoword accuracy was significantly impaired during lucid REM sleep, with lower accuracy compared to wakefulness (median*_diff_* = −1.607, 95% HDI [−2.087, −1.129]) and N1 sleep (median*_diff_* = −0.549, 95% HDI [−1.030, −0.059]). These findings suggest that while participants retained access to words, their ability to process pseudowords was selective impaired in lucid REM sleep.

DDM revealed that lexical decision-making in lucid REM sleep was characterized by reduced evidence accumulation and increased decision caution. Drift rates were significantly lower for both words and pseudowords, indicating diminished processing efficiency (words: median*_diff_* = −0.349, 95% HDI [−0.660, −0.023]; pseudowords: median*_diff_* = −0.806, 95% HDI [−1.072, −0.544], Figure 5B, 5D) compared to wakefulness. A similar reduction in drift rate was observed when comparing lucid REM sleep to N1 sleep, with lower values for both words (median*_diff_* = −0.372, 95% HDI [−0.754, −0.014]) and pseudowords (median*_diff_* = −0.341, 95% HDI [−0.635, −0.056], Figure 5B, 5D).

**Figure 5.**
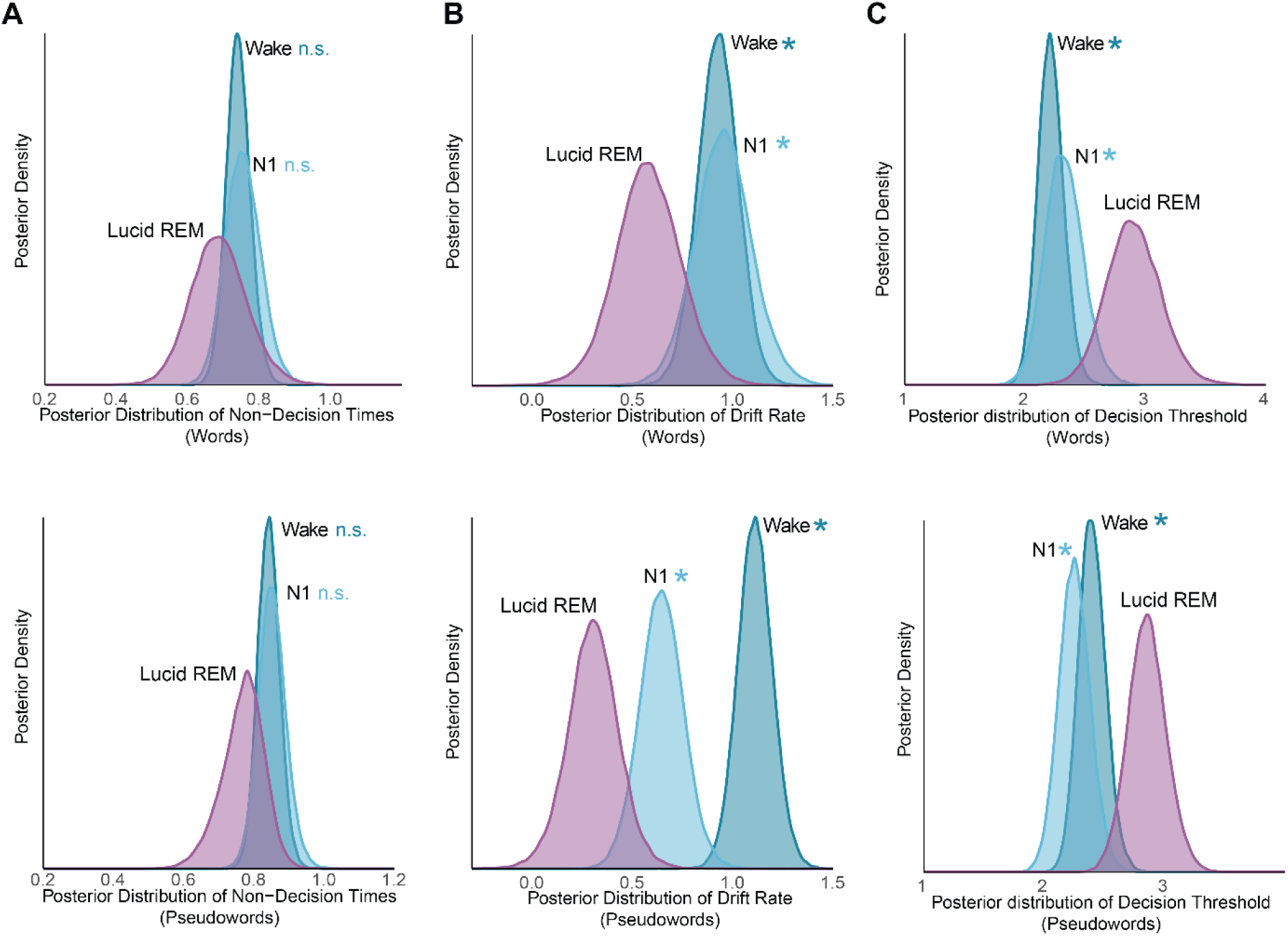
Slower and less efficient lexical decisions during lucid REM sleep. Posterior distributions of decision parameters for words and pseudowords in the NP group, comparing lucid REM sleep, wakefulness, and N1 sleep. Parameters include (A) non-decision time, (B) drift rate, and (C) decision threshold. Here, we specifically compare lucid REM sleep with wakefulness and N1 sleep, while results for N1 vs. wakefulness are reported in the Results section. Asterisks (*) denote significant differences, where the posterior distribution contrast does not overlap with 0. "n.s." indicates non-significant differences, where the posterior distribution contrast overlaps with 0.

In addition to slower evidence accumulation, participants exhibited elevated decision thresholds for both words and pseudowords, suggesting that participants adopted a more cautious decision-making strategy, likely as compensation for reduced processing efficiency. Thresholds were significantly higher in lucid REM sleep compared to wakefulness (words: median*_diff_* = 0.684, 95% HDI [0.259, 1.149]; pseudowords: median*_diff_* = 0.453, 95% HDI [0.102, 0.775], Figure 5C, 5F) and N1 sleep (words: median*_diff_* = 0.578, 95% HDI [0.067, 1.075]; pseudowords: median*_diff_* = 0.599, 95% HDI [0.261, 0.949], Figure 5C, 5F). Despite these changes, non-decision times remained stable across wakefulness, N1 sleep, and lucid REM sleep (all 95% HDIs overlapped with 0, Figure 5A, 5D), indicating that stimulus encoding and response preparation were preserved across states.

Together, these findings reveal a progressive shift in lexical decision-making across wakefulness, N1 sleep, and lucid REM sleep. While N1 sleep preserved word judgment but impaired pseudoword processing, lucid REM sleep was associated with a generalized reduction in processing efficiency. Participants compensated for this inefficiency by adopting a more cautious decision-making strategy, reflected in elevated decision thresholds. Importantly, non-decision times remained stable across all states, suggesting that stimulus encoding and motor preparation were preserved despite changes in decision-making efficiency.

## Discussion

This study investigated how lexical decision-making processes adapt across distinct states of consciousness—wakefulness, N1 sleep, and lucid REM sleep—by integrating behavioural data and computational modelling with the drift diffusion model. Analyzing key decision parameters—non-decision times, drift rates, and decision thresholds—revealed state-specific adaptations in decision-making and how they dynamically reconfigure across states of consciousness. Our findings indicate that N1 sleep preserves both non-decision processes (sensory encoding and motor preparation) and evidence accumulation (drift rate) to support lexical decisions, whereas lucid REM sleep relies primarily on evidence accumulation. Notably, decision-making exhibited a progressive shift across states: while N1 sleep maintained efficient word judgments, it showed selective impairments in pseudoword judgments, reflected in prolonged non-decision times and slower drift rate. In contrast, lucid REM sleep exhibited a generalized decline in decision-making efficiency, characterized by reduced drift rates and elevated decision thresholds for both words and pseudowords. These results suggest that the brain dynamically reallocates cognitive resources to sustain task performance under the distinct state of consciousness.

Although wakefulness served as our baseline, its lexical decision-making profile revealed notable deviations from prior findings, particularly regarding the mechanisms underlying the word advantage. While past research suggests that both drift rate and non-decision time contribute to faster word responses (Donkin et al., 2009; Ratcliff et al., 2004; Wagenmakers et al., 2008), our results indicate that this advantage in wakefulness was driven solely by non-decision time, with no drift rate differences between words and pseudowords. This discrepancy may stem from the spoken auditory lexical decision task, as auditory word recognition relies more on incremental phonological encoding and feedforward activation rather than rapid evidence accumulation (Hickok & Poeppel, 2007; Marslen-Wilson, 1987). Unlike visual word recognition, which engages orthographic feedback and lexical competition (Coltheart et al., 2001; Wagenmakers et al., 2008), spoken word processing unfolds over time, potentially diminishing the role of drift rate. Additionally, NPs exhibited slower response times, reduced accuracy, and lower drift rates, consistent with prior evidence that orexinergic dysfunction disrupts wake-state stability and cognitive vigilance (Dauvilliers et al., 2007; Scammell, 2015). These wakefulness-specific differences underscore the importance of considering baseline cognitive variability when interpreting sleep-related effects.

The ability to perform lexical decisions during N1 sleep indicates that the transition from wakefulness to light sleep does not lead to a complete shutdown of higher-order cognition. Instead, the brain retains functional connectivity and sufficient computational capacity to process external stimuli, particularly those with strong semantic or lexical associations (Andrillon et al., 2016; Kouider et al., 2014; Siclari & Tononi, 2017). Both HPs and NPs preserved the word advantage, characterized by shorter non-decision times and higher drift rates for words. This suggests that familiar, meaningful stimuli continue to benefit from efficient sensory encoding and robust evidence accumulation even in early sleep states (Andrillon & Kouider, 2020; Perrin et al., 1999; Portas et al., 2000). These findings align with theories proposing that N1 sleep retains partial access to the global workspace, allowing meaningful stimuli to penetrate higher-order cognitive systems (Dehaene & Changeux, 2011).

Lucid REM sleep exhibited a distinct lexical decision-making profile, fundamentally diverging from both wakefulness and N1 sleep. Unlike N1 sleep, where both early-stage processing (non-decision time) and evidence accumulation (drift rate) contributed to lexical decision, decision-making in lucid REM sleep relied solely on evidence accumulation. The absence of non-decision time modulation suggests that sensory encoding and motor preparation are no longer limiting factors, likely reflecting the altered neural dynamics of lucidity (Dresler et al., 2015; Voss et al., 2014). Lucid dreaming is associated with increased prefrontal activity and enhanced metacognition, which may enable participants to engage in goal-directed tasks despite the atypical sensory environment (Filevich et al., 2015). In this state, external stimuli may be processed less efficiently, forcing lexical decisions to depend entirely on post-sensory evidence accumulation. These findings suggest that lucidity induces a computational reconfiguration of decision-making, demonstrating how cognitive processes can adapt to hybrid states of consciousness characterized by partial reinstatement of executive control within a modified sensory landscape.

Having established state-specific effects, we also compared lexical decision-making between wakefulness and N1 sleep to examine how the sleeping brain adapts decision strategies. Despite overall slowing in N1 sleep, lexical decisions for words remained relatively preserved, likely due to the robustness of semantic memory networks, which enable rapid access to familiar word representations with minimal cognitive effort (Andrillon et al., 2016; Hickok & Poeppel, 2007). Automatic retrieval of well-established meanings requires minimal cognitive control, making it more resistant to sleep-related impairments. In contrast, lexical decisions for pseudowords were selectively impaired, as reflected in slower response times, lower accuracy, prolonged non-decision times, and reduced evidence accumulation. Unlike words, pseudowords lack semantic associations and require greater bottom-up sensory encoding and cognitive flexibility, which are particularly vulnerable to the neurophysiological constraints of sleep (Andrillon & Kouider, 2020). The prolonged non-decision times observed for pseudowords further suggest inefficiencies in early-stage processing. This pattern reveals a hierarchical prioritization of cognitive resources, where the brain maintains efficient processing for familiar, meaningful stimuli while deprioritizing resource-intensive operations.

Lexical decision-making in lucid REM sleep among narcolepsy participants was marked by slower evidence accumulation (drift rate) and elevated decision thresholds compared to wakefulness, reflecting the cognitive demands of this altered state. Narcolepsy is characterized by intrusive REM features, including heightened internal imagery, dream-like mentation, and dysregulated arousal states (Dauvilliers et al., 2007; Voss et al., 2014), which likely compete with external stimuli for cognitive resources, thereby introducing cognitive noise and reducing the efficiency of evidence accumulation. To compensate for this increased uncertainty, participants adopted a more cautious decision strategy, setting higher decision thresholds to mitigate errors (Forstmann et al., 2016; Ratcliff et al., 2016). Despite these adjustments, non-decision times remained stable, indicating that sensory encoding and motor preparation were preserved, even as evidence accumulation became less efficient.

Unlike N1 and lucid REM sleep, participants exhibited no evidence of lexical decision-making during N2 or non-lucid REM sleep in either group. Behavioural and computational analyses revealed no significant differences between words and pseudowords, suggesting that higher-order cognitive functions and evidence accumulation were absent in these states. This aligns with research indicating that deeper sleep states, such as N2 and REM, involve reduced thalamocortical connectivity and sensory disconnection, restricting processing to lower-level sensory areas with limited capacity for decision-related computations (Andrillon & Kouider, 2020; Nir et al., 2011). However, this finding contrasts with Türker et al. (2023), who reported above-chance responses to verbal stimuli across all sleep states, including N2 and REM. One explanation for this discrepancy is that while participants in Türker et al. (2023) may have exhibited stimulus-driven motor responses, our drift diffusion modelling suggests that these responses were not indicative of true lexical decision-making. Instead, they likely reflect automatic sensory-motor processing, such as priming effects or residual auditory-motor coupling, which can persist in deep sleep despite the absence of volitional decision processes (Andrillon et al., 2016; Kouider et al., 2014). This interpretation reinforces the idea that higher-order linguistic operations require a minimal level of wake-like cortical integration, which may be present in N1 and lucid REM but absent in N2 and non-lucid REM sleep.

By integrating behavioural and computational approaches, this study provides a new framework for studying high-order cognition beyond wakefulness, offering insights into how the brain maintains residual cognitive functions in altered states of consciousness. While our Bayesian and DDM analyses accounted for unbalanced trial conditions, future studies with larger sample sizes and overnight paradigms could further enhance statistical power (Ratcliff et al., 2016). Additionally, our investigation of lucid REM sleep was limited to participants with narcolepsy, raising questions about the generalizability of these effects to healthy individuals (Baird et al., 2019). Future studies using high-density EEG or neuroimaging could further elucidate the neural mechanisms underlying sleep-based decision-making. These findings have broad implications for sleep’s role in cognition, the nature of conscious processing across vigilance states, and the neural mechanisms that shape decision-making under varying levels of arousal and awareness.

## Methods

This study presents a novel analysis of data originally collected in 2020 at the Sleep Clinic of Pitié-Salpêtrière Hospital, France, as part of an experiment previously published (Türker et al., 2023). The study adhered to the Declaration of Helsinki, and ethical approval was granted by the local ethics committee (CPP Ile-de-France 8). All participants provided written informed consent before participation.

Thirty individuals diagnosed with narcolepsy (NP; 14 women; mean age: 35 ± 11 years) and 22 healthy participants (HP; 10 women; mean age: 24 ± 4 years) were recruited. Participants with narcolepsy were diagnosed according to international diagnostic criteria and recruited from the National Reference Center for Narcolepsy at the Pitié-Salpêtrière Hospital. Among NPs, 80% reported frequent lucid dreaming (≥3 lucid dreams per week), whereas none of the HPs reported a history of lucid dreaming. Three participants (two NPs and one HP) were excluded due to technical issues during data acquisition, leaving 27 NPs (21 frequent lucid dreamers) and 21 HPs in the final analyses. Participants were compensated financially for their involvement. For detailed demographic and clinical information, see the previously published study (Türker et al., 2023).

## Experimental design

### Task Overview

Participants performed a lexical decision task in which they determined whether auditory stimuli were real words or pseudowords. Responses were indicated via brief contractions of facial muscles: the corrugator (frowning) and zygomatic (smiling) muscles. The muscle-response mappings were counterbalanced across participants. Stimuli were presented in pseudorandomized order, ensuring each stimulus was presented only once to prevent repetition effects. Participants completed a 10-minute familiarization session before data collection to practice the task and ensure comfort with the auditory stimuli, which were played at an average volume of 48 dB and adjusted for individual audibility.

### Nap Protocol

Participants with narcolepsy completed five 20-minute nap sessions, interspersed with 80-minute breaks, while HPs completed a single uninterrupted 100-minute daytime nap. Each nap session consisted of 10 active ("ON") periods, during which six stimuli (three words, three pseudowords) were presented every 9–11 seconds against a background of continuous white noise. These "ON" periods alternated with 1-minute "OFF" intervals, during which only white noise was delivered. Across sessions, 60 stimuli (30 words, 30 pseudowords) were presented per participant, with presentation lists randomized to mitigate order effects.

### Stimuli

Auditory stimuli were selected from the MEGALEX database (Ferrand et al., 2018) and included French words and pseudowords spoken by a female voice. Stimuli were standardized to a duration of 690 ms and controlled for frequency and emotional valence. To ensure consistency, each participant received five unique stimulus lists, randomized across nap sessions. Stimuli were delivered via speakers using Psychtoolbox in MATLAB (MathWorks), with a randomized inter-stimulus interval of 9–11 seconds.

### Electrophysiological recording

Electrophysiological data were collected using a 10-channel EEG setup (Fp1, Fp2, Cz, C3, C4, Pz, P3, P4, O1, O2), following the international 10–20 system. Signals were referenced to the right mastoid (A2 electrode). Additional recordings included electrooculography (EOG) from two electrodes positioned to capture eye movements, electromyography (EMG) from three channels (chin muscles for sleep staging and the zygomatic and corrugator muscles to record behavioral responses), and electrocardiography (ECG) from one channel to record heart activity. All signals were recorded continuously at a sampling rate of 2,048 Hz using a Grael 4K PSG/EEG amplifier (Medical Data Technology, Compumedics).

### Sleep scoring and identification of lucid dream

Sleep stages were scored offline by a certified sleep expert according to American Academy of Sleep Medicine guidelines (Berry et al., 2017) using Profusion (Compumedics). EEG and EOG signals were filtered between 0.3–15 Hz, EMG between 10–100 Hz, and ECG between 0.3–70 Hz. Sleep stages were scored in 30-second epochs as wakefulness, N1, N2, N3, or REM sleep. Micro-arousals were defined as alpha activity lasting 3–15 seconds, with arousals exceeding 15 seconds classified as wakefulness. For REM sleep, micro-arousals were further characterized by transient increases in EMG tone. Trials containing micro-arousals were excluded from subsequent analyses.

Lucid REM sleep was identified based on participants’ self-reports following each nap session. If a participant reported a lucid dream, all REM epochs from that session were classified as lucid REM sleep. No HPs reported lucid dreams.

### Muscle Response Analysis

EMG signals were segmented into 10-second mini-epochs corresponding to sleep stages. Mini-epochs with micro-arousals were excluded. Muscle contractions were classified as valid responses if at least two consecutive contractions were detected; single contractions (twitches) were excluded as non-responses. Scoring reliability was validated by reanalyzing 10% of the data with a second blinded scorer, yielding 84% agreement.

### Drift diffusion model analysis

To investigate the mechanisms underlying lexical decision-making across wakefulness and sleep stages, we employed the Drift Diffusion Model (DDM), a well-established framework for modeling two-choice decision-making tasks (Ratcliff et al., 2016). The DDM assumes that decisions arise from a continuous process of evidence accumulation, where sensory information about the two options (e.g., words and pseudowords) is integrated over time until a decision threshold is reached. The model decomposes behavioral data (accuracy and response times) into distinct cognitive parameters, providing insights into the underlying decision-making processes. The DDM decomposes this process into four main components: the starting point (***z***), which indicates a predecision bias; the nondecision time (***t***), which covers factors unrelated to the actual decision and is often linked to the encoding, motor execution, and lexical access in lexical tasks; the drift rate (***v***), which represents the speed of information accumulation; and the decision threshold (***a***), which indicate when enough evidence has been collected to make a decision.

Previous findings suggest that differences in lexical decision behavior are primarily driven by drift rate and non-decision time (Donkin et al., 2009; Ratcliff et al., 2004). However, to account for potential variations in decision-making strategies across wakefulness and sleep, we also included decision threshold (a) in our model. This approach allowed us to examine how drift rate (v), non-decision time (t), and decision threshold (a) were modulated by word type and state of consciousness (wakefulness and sleep states). Predecision bias (starting point) was estimated at the participant level and assumed to remain constant across word type and sleep states, as our primary interest lay in the interaction between evidence accumulation, decision thresholds, and state-dependent cognitive dynamics (Donkin et al., 2009; Herz et al., 2022; Ratcliff et al., 2004). Separate HDDM analyses were conducted for HP and NP to assess group-specific effects (Figure 1C).

We used a Bayesian hierarchical approach with the HDDM 0.8 tool in Docker to estimate these parameters (Pan et al., 2022), assuming that participants’ parameters are drawn from a shared distribution. We applied Markov Chain Monte Carlo (MCMC) sampling to generate 10,000 samples, discarding the first 1,000 as burn-in. Model convergence was checked by inspecting trace plots, autocorrelation, and the Gelman–Rubin R-hat statistic (ensuring R-hat < 1.1). We applied HDDM regression analysis separately for HP and NP using the following model:

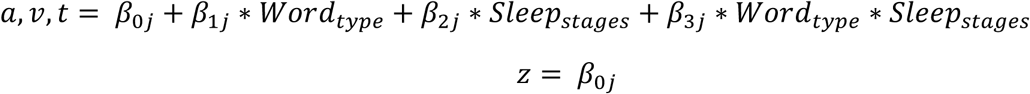

### Statistics

We only included response trials where stimuli were presented in the current study. Additionally, trials with microarousal (HP: 15%; NP: 13.6%) were excluded from the analysis. N3 (0.07%) and REM (1%) sleep trials in HP were excluded due to insufficient trial numbers. To eliminate the possibility of random muscle contractions, we excluded trials in which participants exhibited only a single muscle contraction (HP: 0.6%; NP: 1.5%). Lastly, responses with times less than 0.69 seconds or greater than 9.9 seconds were excluded from the analysis. Moreover, outliers were excluded according to the conservative criterion of mean ± 2.5 median absolute deviation (MAD) based on RT.

Our study aimed to uncover the computational processes underlying lexical decision-making across wakefulness and sleep in healthy participants (HP) and individuals with narcolepsy (NP) using the drift diffusion framework. Participants performed a lexical decision task (LDT) during wakefulness and continued the task throughout sleep, using facial muscle contractions (zygomatic and corrugator muscles) to indicate whether spoken stimuli were words or pseudowords in the French lexicon (Figure 1A). Polysomnography, along with additional EMG sensors, was used to confirm sleep stages and capture behavioral responses. This setup allowed us to collect both accuracy and RT data for lexical judgments during wakefulness and sleep. To analyze behavioral performance, we applied a Bayesian linear mixed model (BLMM) to trial-level accuracy and RT data. Fixed factors included word type (words vs. pseudowords) and sleep stages, while participants were modeled as a random factor (Figure 2). BLMM is particularly suited for analyzing hierarchical data structures, addressing individual variability and imbalances in trial numbers (Franke & Roettger, 2019; Gelman et al., 2014; Sorensen et al., 2016).

The model is specified as follows:

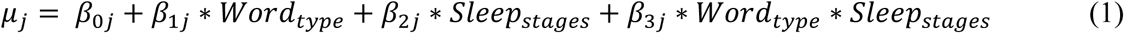

Where *μ*_*j*_ represents either response accuracy or reaction time, and *j* represents the subject. For response accuracy, we used the Bernoulli family to model the binary data. For reaction time, we employed the shifted lognormal family to appropriately model the RT data.

During the analysis of reaction time, we conducted a control analysis by adding response accuracy as a fixed factor to examine whether correct or incorrect responses would show different reaction times:

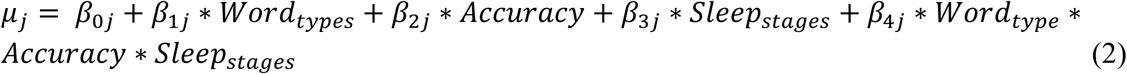

For each model, we ran four MCMC chains with 5,000 samples each, discarding the first 500 samples as a warm-up. We assessed model convergence using the Gelman–Rubin R-hat statistic, ensuring (R-hat < 1.1). Statistical inferences were based on the 95% Highest Density Interval (HDI) of the posterior distribution. Effects were considered significant if the 95% HDI did not include 0.

## Data availability

All data will be available on the Open Science Framework (OSF) upon publication: https://osf.io/r8szg/

## Code availability

All analysis codes will be accessible on the Open Science Framework (OSF) upon publication: https://osf.io/r8szg/

## Acknowledgements

We would like to express our gratitude to Dr. Menglu Chen for offering valuable feedback and suggestions on this article and its visualizations. The research was supported by the Ministry of Science and Technology of China STI2030-Major Projects (No. 2022ZD0214100), National Natural Science Foundation of China (No. 32171056), General Research Fund (No. 17614922) of Hong Kong Research Grants Council to X. H. The funders had no involvement in the study design, data collection and analysis, publication decisions, or manuscript preparation.

## Contributions

T.X. conceived and designed the project. T.X. performed the formal data analysis. T.X. and X.H wrote the original draft. C.H., B.T., E.M., L.N., I.A., D.O., and X.H reviewed and edited the article, providing valuable suggestions.

---

**Table S1.** Mean and SEM of trial numbers across sleep states and word types in HPs group.

| | Wake (Mean ± SE) | Healthy Group<br>N1(Mean ± SE) | N2(Mean ± SE) |
| --- | --- | --- | --- |
| Words | 48.2 ± 7.92 | 5 ± 0.89 | 3 ± 0.59 |
| Pseudowords | 46.7 ± 7.67 | 3.33 ± 0.44 | 2.77 ± 0.71 |

**Table S2.** Mean and SEM of trial numbers across sleep states and word types in NPs group.

| | Wake<br>(Mean ± SE) | Individual with narcolepsy<br>N1<br>(Mean ± SE) | N2<br>(Mean ± SE) | REM<br>(Mean ± SE) | Lucid REM<br>(Mean ± SE) |
| --- | --- | --- | --- | --- | --- |
| Words | 26.4 ± 5.79 | 9.04 ± 1.68 | 8.92 ± 9.17 | 12.2 ± 3.97 | 14.4 ± 2.15 |
| Pseudowords | 27.1 ± 5.94 | 8.69 ± 1.53 | 9.17 ± 1.78 | 11.6 ± 3.64 | 15.5 ± 2.24 |

**Figure S1.**
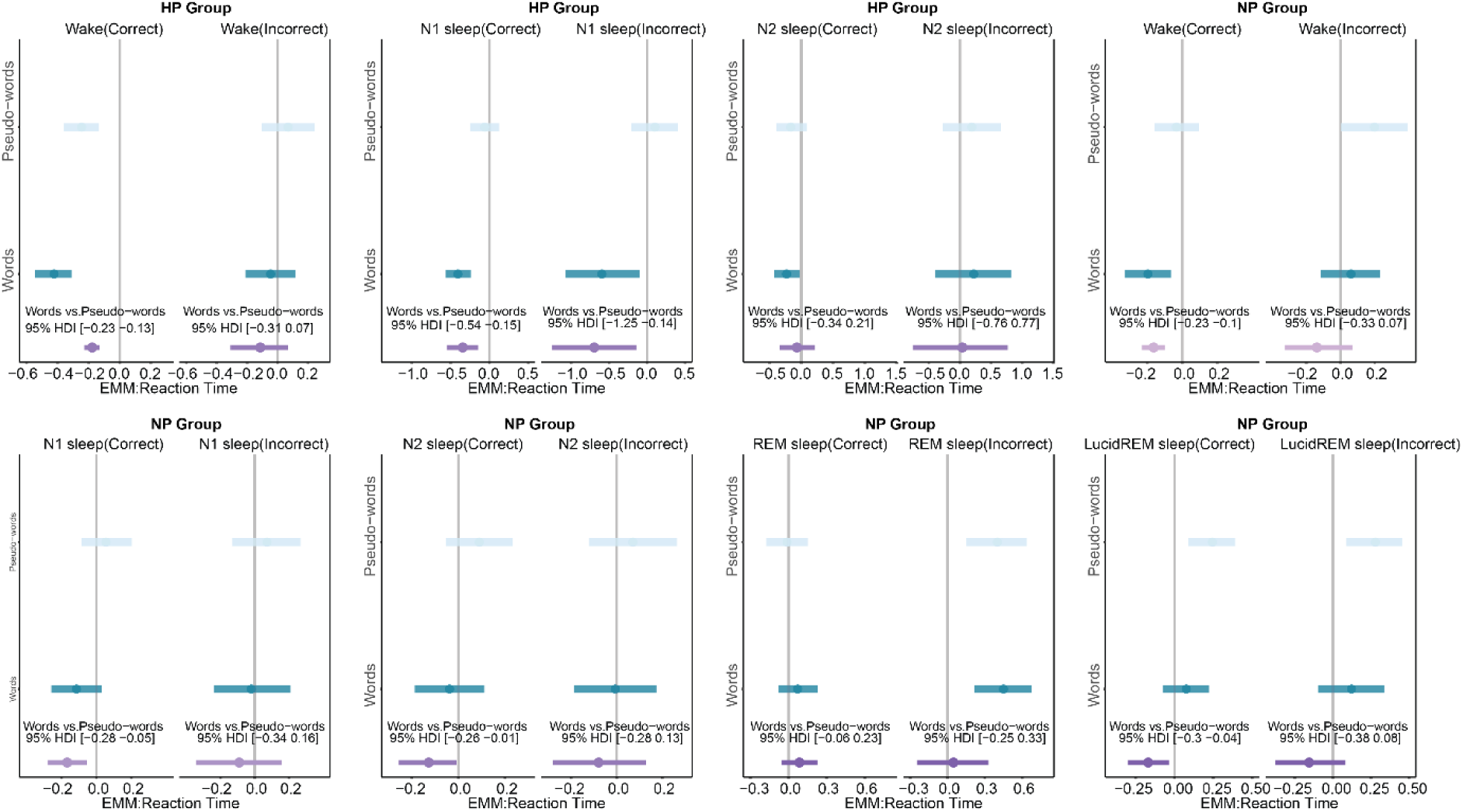
Differences in Reaction Time between Words and Pseudowords across wake/sleep stages in Healthy Participants (HP) and Participants with Narcolepsy (NP). This figure depicts the reaction time differences for words and pseudowords, considering both correct and incorrect responses. The X-axis denotes the estimated mean of reaction time. The light blue line represents the 95% Highest Density Interval (HDI) of the posterior probability for pseudowords, while the green line represents words. The purple line indicates the contrast between words and pseudowords. If the purple line overlaps with the 0 (gray line), it suggests no significant difference between words and pseudowords. Conversely, if the purple line does not overlap with 0, it indicates a significant difference between words and pseudowords.

**Figure S2.**
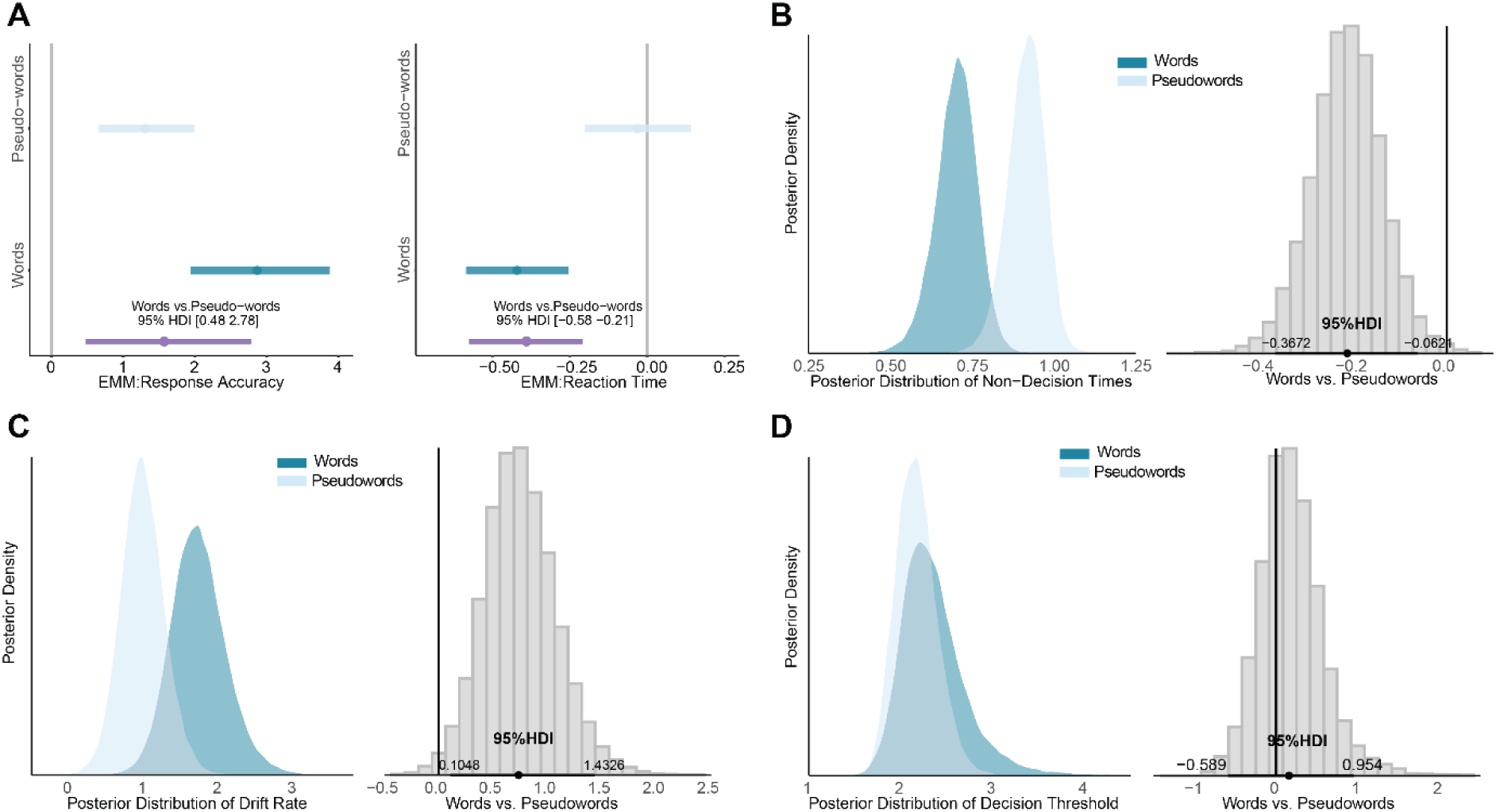
Mechanism of lexical decision during N1 in healthy participants. (A)BLMM results for response accuracy and reaction time in lexical decisions during N1 sleep in healthy participants. The X-axis represents the estimated mean response accuracy or reaction time. The light blue line indicates the 95% Highest Density Interval (HDI) of the posterior probability for pseudowords, while the green line represents words. The purple line denotes the contrast between words and pseudowords. If the purple line overlaps with 0 (gray vertical line), there is no significant difference between words and pseudowords. If the purple line does not overlap with 0, it indicates a significant difference between words and pseudowords. (B, C, D) Posterior distributions of non-decision times, drift rate, and decision threshold for words and pseudowords during N1 and lucid REM sleep, along with their contrasts. The left panels show the fitted posterior distributions for words and pseudowords. The right-side histogram plots display the contrasts between words and pseudowords, with horizontal black lines representing the 95% HDI and vertical gray lines denoting 0. If 0 is not within the 95% HDI, the difference is considered statistically significant.

**Figure S3.**
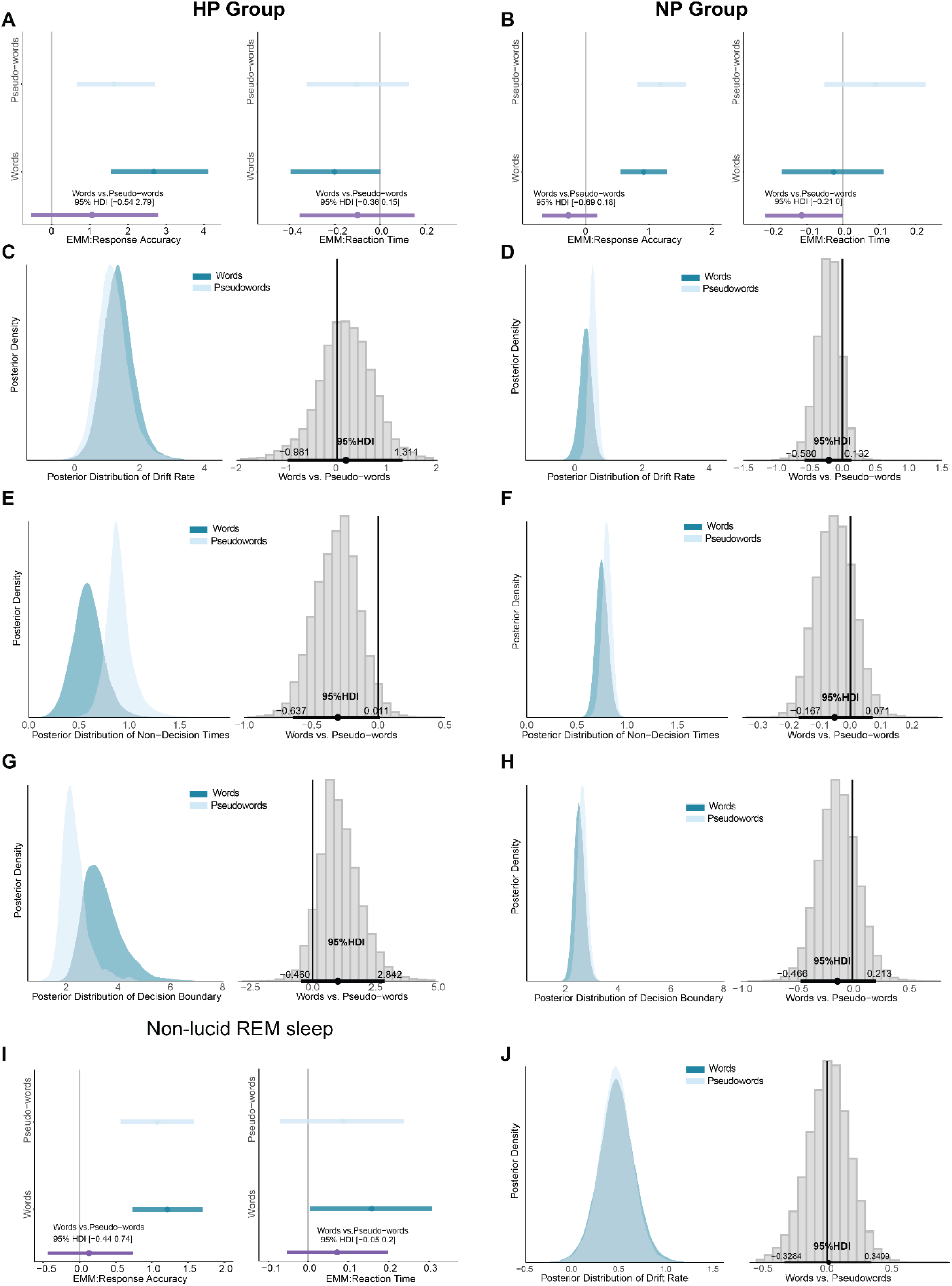
Absence of lexical decision-making during N2 sleep (HP & NP) and non-lucid REM sleep (NP) (A, B, I) Response accuracy and reaction time during N2 and non-lucid REM sleep. The X-axis shows mean accuracy and reaction time. The light blue and green lines represent pseudowords and words, respectively. The purple line indicates the difference between them. If the purple line overlaps with 0 (gray vertical line), there is no significant difference. (B, C, D, E, F, G, H, J) Distributions of non-decision time, drift rate, and decision threshold. Left panels show results for words and pseudowords. Right panels display their differences, with black lines marking the 95% HDI and the gray vertical line indicating 0. If 0 is within the 95% HDI, there is no significant difference, confirming that lexical decision-making is absent in these states.

